# Dissociated hippocampal neurons exhibit distinct Zn^2+^ dynamics in a stimulation method-dependent manner

**DOI:** 10.1101/2020.01.03.894501

**Authors:** Lynn Sanford, Amy E. Palmer

## Abstract

Ionic Zn^2+^ has increasingly been recognized as an important neurotransmitter and signaling ion in glutamatergic neuron pathways. Intracellular Zn^2+^ transiently increases as a result of neuronal excitation, and this Zn^2+^ signal is essential for neuron plasticity, but the source and regulation of the signal is still unclear. In this study we rigorously quantified Zn^2+^, Ca^2+^ and pH dynamics in dissociated mouse hippocampal neurons stimulated with bath application of high KCl or glutamate. While both stimulation methods yielded Zn^2+^ signals, Ca^2+^ influx, and acidification, glutamate stimulation induced more sustained high intracellular Ca^2+^ and a larger increase in intracellular Zn^2+^. However, the stimulation-induced pH change was similar between conditions, indicating that a different cellular change is responsible for the stimulation-dependent difference in Zn^2+^ signal. This work provides the first robust quantification of Zn^2+^ dynamics in neurons using different methods of stimulation.

---

Zn^2+^ is an essential metal ion cofactor that regulates the structure or function of thousands of mammalian proteins^1,2^. Zn^2+^ has also been shown to be integral to signaling in specific cellular systems, including in a subset of glutamatergic neurons throughout different brain regions^3^. In these neurons, Zn^2+^ localizes within synaptic vesicles and is released into the synapse along with glutamate upon stimulation^4–9^. It interacts allosterically or competitively with a number of postsynaptic neurotransmitter receptors, most notably NMDA-type glutamate receptors, to modulate synaptic potentiation^10–14^. Intracellular labile Zn^2+^ also increases in stimulated hippocampal neurons and is known to be important for synaptic growth and plasticity^15–21^.

In dissociated hippocampal neuron culture, stimulation with glutamate/glycine^22^ or KCl^23^ has been shown to increase intracellular Zn^2+^, and this Zn^2+^ signal has important downstream signaling consequences. This Zn^2+^ has been suggested to arise from an intracellular source in a Ca^2+^-dependent manner, and previous studies have indicated that during glutamate stimulation the Zn^2+^ signal may be downstream of Ca^2+^-induced neuronal acidification^22,24^. However, no study to this point has compared Zn^2+^ responses in dissociated neurons using different stimulation methods, which could further clarify the mechanism of Zn^2+^ mobilization. In this study we examined Zn^2+^ signals generated during either glutamate or KCl stimulation of dissociated mouse hippocampal neuron cultures through fluorescence imaging of Zn^2+^, Ca^2+^ and pH. We found that different stimulation methods generated different intracellular Zn^2+^ and Ca^2+^ dynamics, but these differences were independent of observed pH changes, implicating an additional process in stimulation-dependent intracellular Zn^2+^ mobilization.

## RESULTS AND DISCUSSION

### Intracellular Zn^2+^ increases to varying extents upon different neuronal culture stimulations

To determine how intracellular Zn^2+^ dynamics differ in dissociated hippocampal neurons depending on stimulation method, we applied 50 mM KCl or 50 μM glutamate to neuron cultures and imaged Zn^2+^ with FluoZin-3 AM (Figure 1). Both of these stimulations resulted in elevated intracellular Zn^2+^, with Zn^2+^ levels approximately reverting to baseline levels after stimulation washout (Figure 1A, left panels). Generally, glutamate stimulation gave rise to ~2-fold larger peak intracellular Zn^2+^ increases than those observed upon KCl stimulation in the same stimulation time period (Figure 1B, Table 1, Table 2). Furthermore, Zn^2+^ signals evoked by both KCl and glutamate stimulation could be intensified by the addition of exogenous Zn^2+^ (Figure 1A, right panels), and in this case the peak Zn^2+^ levels were similar regardless of stimulation method (Figure 1B, Table 1, Table 2). Neurons thus have comparable permeability to extracellular Zn^2+^ during KCl and glutamate stimulation, although permeability may be somewhat higher during KCl stimulation given the lower intracellular Zn^2+^ signal in endogenous conditions.

**Table 1.**
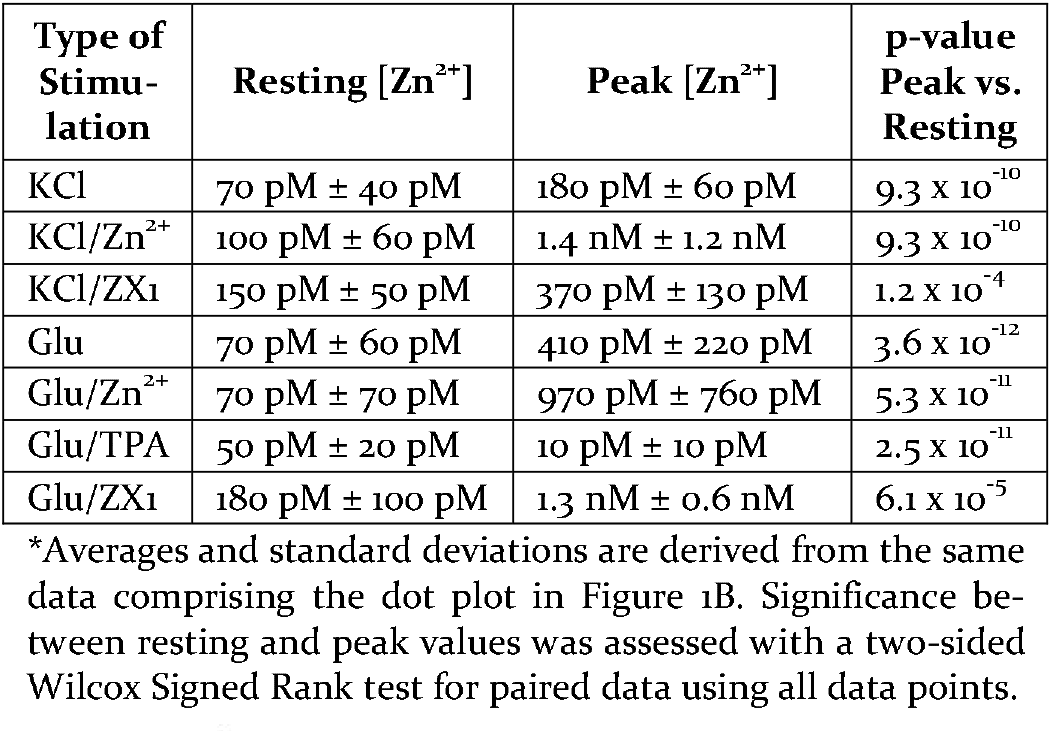
Approximate Zn^2+^ concentrations in different stimulation conditions, from FluoZin-3 AM data*.

**Table 2.** Results of significance tests pairwise between stimulation conditions for FluoZin-3 AM, Fluo-4 AM, and BCECF AM data*.

| Dye/imaging phase comparison | KCl<br>vs.<br>KCl/Zn<sup>2+</sup> | KCl<br>vs.<br>Glu | Glu<br>vs.<br>Glu/Zn<sup>2+</sup> | Glu<br>vs.<br>Glu/TPA | KCl/Zn<sup>2+</sup><br>vs.<br>Glu/Zn<sup>2+</sup> | KCl/ZX1<br>vs.<br>Glu/ZX1 |
| --- | --- | --- | --- | --- | --- | --- |
| Maximum excitation FluoZin-3 FS | $8.4 \times 10^{-16}$ | $2.2 \times 10^{-7}$ | $3.1 \times 10^{-9}$ | $< 2.2 \times 10^{-16}$ | 0.355 | $8.6 \times 10^{-7}$ |
| Recovery FluoZin-3 FS post-excitation | $< 2.2 \times 10^{-16}$ | 0.592 | $1.1 \times 10^{-5}$ | $6.6 \times 10^{-11}$ | $2.4 \times 10^{-7}$ | 0.013 |
| Maximum excitation Fluo-4 F/F<sub>max</sub> | 0.858 | 0.011 | $1.9 \times 10^{-5}$ | | 0.016 | |
| Summed excitation Fluo-4 F/F<sub>max</sub> | 0.858 | $6.4 \times 10^{-5}$ | $3.7 \times 10^{-6}$ | | 0.109 | |
| Minimum excitation pH (BCECF) | $4.1 \times 10^{-5}$ | 0.050 | 0.017 | 0.135 | 0.010 | |
| Recovery pH post-excitation | $7.3 \times 10^{-4}$ | $1.2 \times 10^{-8}$ | 0.079 | 0.092 | $1.6 \times 10^{-11}$ | |

\*In all cases, significance was assessed with a two-sided Mann-Whitney U test for unpaired data as applied to all data points from each condition. FS=fractional saturation; F/F<sub>max</sub>=fluorescence/maximum fluorescence.

**Figure 1.**
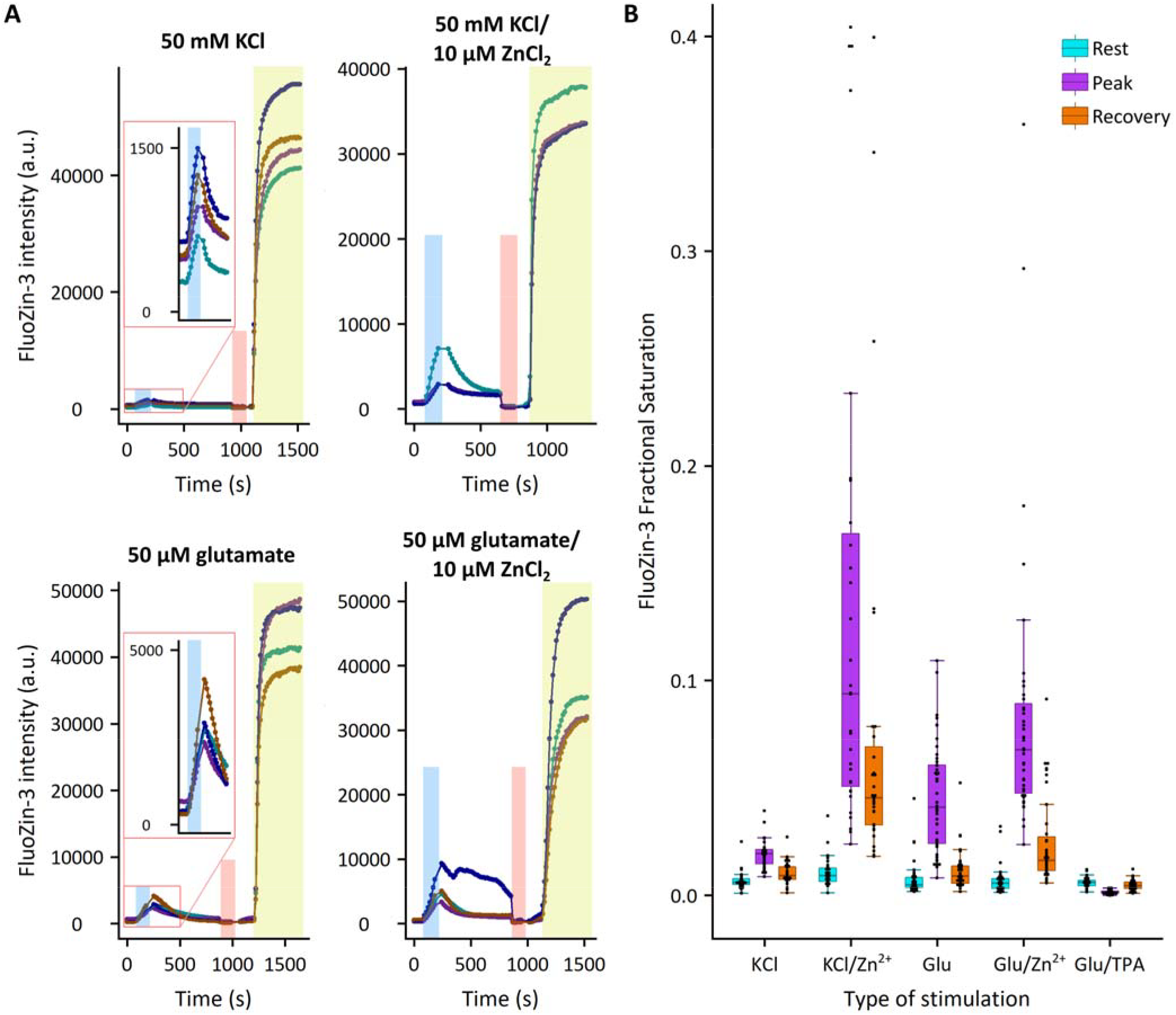
Measurement of stimulation-dependent intracellular Zn^2+^ responses with FluoZin-3 AM. (A) Representative FluoZin-3 intensities upon different stimulations. Each graph is a separate experiment using a different stimulation. Each individual trace represents the average intensity within a different cell in a single field of view. Experiments consisted of stimulation (blue box), followed by a period of recovery before addition of 10 µM TPA (red box) to measure minimum FluoZin-3 signal, then addition of 10 µM ZnCl_2_/0.5 µM pyrithione (yellow box) to achieve maximum FluoZin-3 signal. These minimal and maximal values were used to calculate fractional saturation of the sensor before, during, and after stimulation. (B) Box/dotplot of measured FluoZin-3 fractional saturation (FS) in different stimulation conditions. Each dot represents values obtained from an ROI in a single cell. KCl/Zn^2+^, Glu/Zn^2+^, and Glu/TPA cells were stimulated in the presence of 10 µM ZnCl_2_ or 10 µM TPA. Rest values represent the average FS before stimulation, peak values represent a 3-frame average around the maximal FS obtained during or directly after stimulation, and recovery values represent a 5-frame average FS before addition of TPA for calibration. Sample sizes: KCl = 31 cells from 7 separate experiments; KCl/Zn^2+^ = 31 cells from 9 separate experiments; Glu = 46 cells from 10 separate experiments; Glu/Zn^2+^ = 39 cells from 8 separate experiments; Glu/TPA = 40 cells from 8 separate experiments.

While the observed stimulation-dependent rise in intracellular Zn^2+^ could be related to the potential release of a synaptic Zn^2+^ pool in these neuron cultures, we were previously unable to visualize synaptic Zn^2+^ in dissociated culture^23^. Based on this and other previous work^22^, we suspected that synaptically released Zn^2+^ was not the primary source of the observed intracellular Zn^2+^ signal. We confirmed this hypothesis by imaging stimulation-induced Zn^2+^ responses in the presence of tris(2-pyridylmethyl)amine (TPA), a membrane-permeable Zn^2+^ chelator, or ZX1, a membrane-impermeable Zn^2+^ chelator that has previously been shown to abrogate diffusion of Zn^2+^ across the synaptic cleft^25^ (Figure 2). While incubation of neurons in TPA shortly before and during stimulation completely abolished the Zn^2+^ rise (Figure 1B, Table 1, Table 2), neurons still exhibited a significant intracellular Zn^2+^ increase upon stimulation in the presence of ZX1 (Figure 2, Table 1, Table 2), indicating that the source of labile Zn^2+^ mobilized upon stimulation is primarily intracellular.

**Figure 2.**
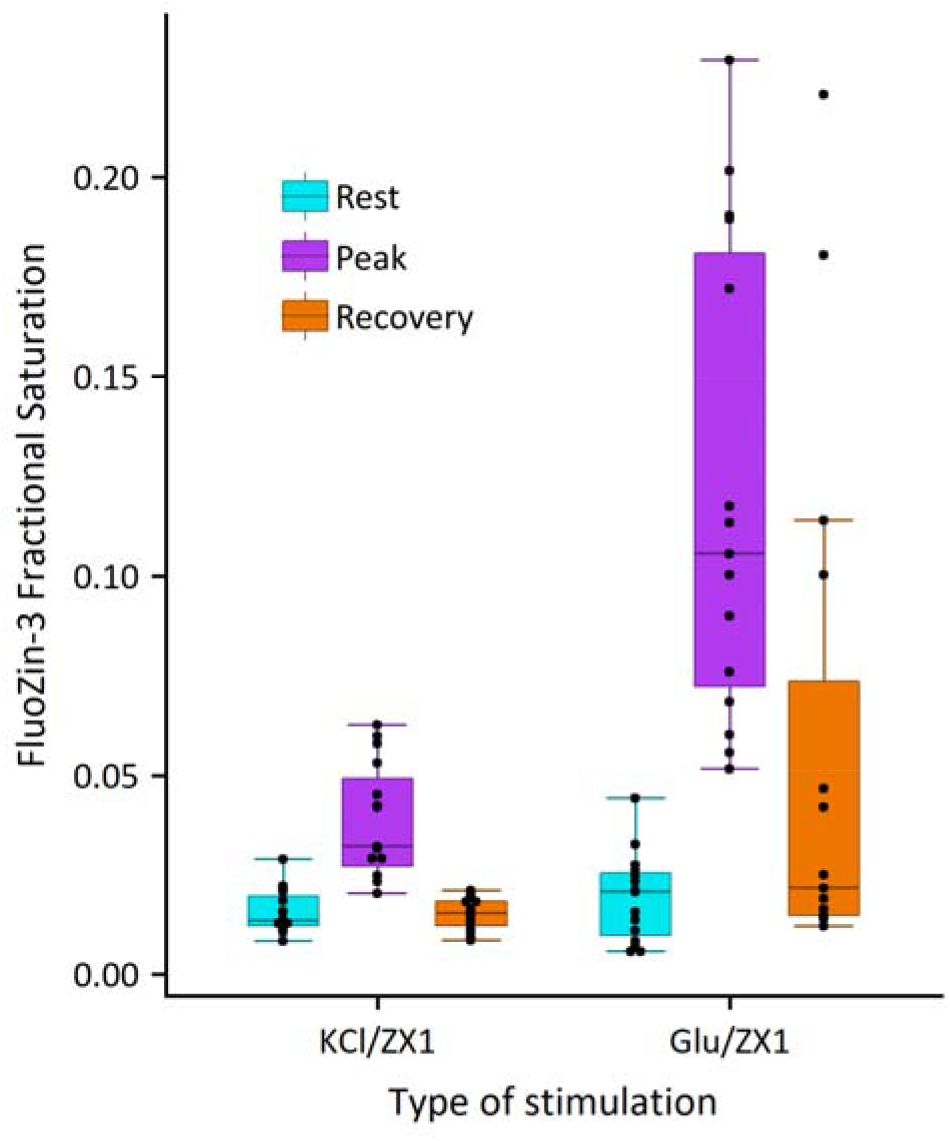
Measurement of stimulation-dependent intracellular Zn^2+^ responses in the presence of 20 µM Zn^2+^ chelator ZX1. Data are shown in a box/dot plot with each dot representing a value obtained from an ROI in a single cell. Rest values represent the average FS before ZX1 addition, peak values represent a 3-frame average around the maximal FS obtained during or directly after stimulation, and recovery values represent a 5-frame average FS before addition of TPA for calibration. Sample sizes: KCl/ZX1 = 15 cells from 2 separate experiments; Glu/ZX1 = 15 cells from 2 separate experiments.

### Sustained calcium influx upon glutamate stimulation may induce a higher Zn^2+^ signal

In order to determine why the endogenous Zn^2+^ signals observed upon KCl or glutamate stimulation differ in magnitude, we first investigated whether intracellular Ca^2+^ dynamics were different by imaging neurons during stimulation with the fluorescent Ca^2+^ dye Fluo-4 AM (Figure 3). We found that while Ca^2+^ significantly increased in all of our stimulation conditions (Figure 3B), Ca^2+^ influx was more sustained over the course of stimulation in glutamate-treated neurons (Figure 3A). Peak Ca^2+^ responses were slightly higher upon glutamate stimulation as compared to KCl stimulation (Figure 3B, Table 2), but this difference became significantly more pronounced if measurements were summed over the course of the two-minute stimulation period (Figure 3C, Table 2). Peak Zn^2+^ concentrations were observed at or after the end of stimulation (Figure 1A), in contrast to the immediate peak Ca^2+^ concentrations upon stimulation onset. The greater Zn^2+^ signals observed upon glutamate stimulation are possibly indicative of a Ca^2+^-dependent process, as the more sustained Ca^2+^ elevation in glutamate-stimulated cells corresponds to a higher Zn^2+^ response.

**Figure 3.**
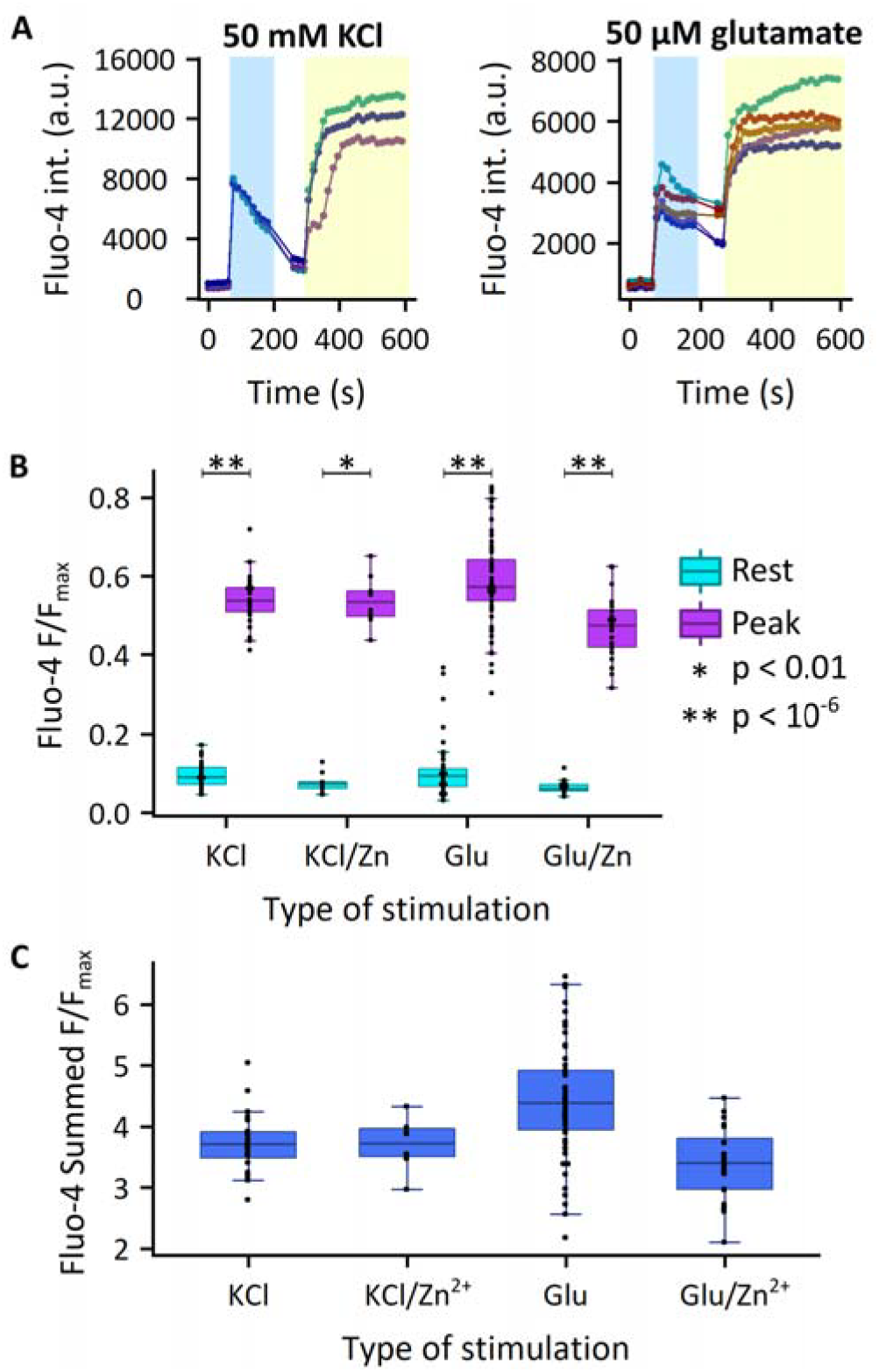
Measurement of stimulation-dependent intracellular Ca^2+^ responses with Fluo-4 AM. (A) Representative Fluo-4 intensities upon stimulation with 50 mM KCl (top) or 50 µM glutamate (bottom). Each individual trace represents the average intensity within a different cell in a single field of view. Experiments consisted of stimulation (blue box), followed by washout and addition of 5 mM CaCl_2_/5 µM ionomycin (yellow box) to achieve maximum Fluo-4 signal. (B) Box/dotplot of normalized Fluo-4 fluorescence in different stimulation conditions. Each dot represents values obtained from an ROI in a single cell. KCl/Zn^2+^ and Glu/Zn^2+^ cells were stimulated in the presence of 10 µM ZnCl_2_. Rest values represent the average F/F_max_ before stimulation, and peak values represent a 3-frame average around the maximal F/F_max_ obtained during stimulation. Sample sizes: KCl = 28 cells from 8 separate experiments; KCl/Zn^2+^ = 10 cells from 2 separate experiments; Glu = 57 cells from 8 separate experiments; Glu/Zn^2+^ = 24 cells from 3 separate experiments. Significance between resting and peak values was assessed with a two-sided Wilcox Signed Rank test for paired data using all data points in each condition (KCl: p=7.5e-9, KCl/Zn^2+^: p=0.002, Glu: p=5.3e-11, Glu/Zn^2+^: p=1.2e-7). (C) Box/dotplot of summed Fluo-4 fluorescence in different stimulation conditions, as determined by summing all F/F_max_ values for the 2 minute stimulation period. Cell ROIs and F/F_max_ values are from the same cells as used in part B.

In the presence of 10 μM exogenous Zn^2+^, Ca^2+^ influx is significantly reduced upon glutamate stimulation, but not during KCl stimulation (Figure 3B, Table 2). This observation is consistent with Zn^2+^ inhibition of glutamate receptors, which has been extensively documented at similar or lower extracellular Zn^2+^ concentrations^10,11^.

### Neurons exhibit different acidification dynamics upon stimulation, but pH does not explain differences in Zn^2+^ signals

There is evidence in the literature that intracellular Zn^2+^ may rise due to Ca^2+^/H^+^ exchange and subsequent acidification of neurons during stimulation, whereby acidification causes release of Zn^2+^ from cytosolic Zn^2+^-binding proteins^22,24^. To determine whether the different magnitudes of Zn^2+^ signals observed upon different stimulation methods were attributable to pH changes, we imaged stimulated neurons with the ratiometric pH-sensing fluorescent dye 2′,7′-bis-(2-carboxyethyl)-5(6)-carboxyfluorescein (BCECF) AM (Figure 4). Comparing fluorescence ratios (Figure 4 A, B) to standard curve ratios measured on the same day of imaging (Figure 4C), we determined that neurons acidified regardless of stimulation method, usually to between pH 6 and pH 7 (Figure 4D). The dynamics of the stimulation-dependent pH change directly mirrors the Zn^2+^ signal dynamics, in which maximal changes were observed either immediately before or immediately after stimulation washout (Figure 1A, Figure 4B). However, the magnitude of acidification was not significantly different between KCl and glutamate stimulation conditions at this timepoint, thus implying that pH is not the primary cause of the difference in Zn^2+^ signals between the two stimulations (Figure 4D, Table 4). We saw further evidence for this conclusion due to our observation that in glutamate-stimulated neurons the intracellular pH remained significantly lower in the “Recovery “ phase 6-7 minutes after stimulation washout (Figure 4D, Table 4), potentially due to the observed sustained intracellular Ca^2+^ response in this condition. In our Zn^2+^ measurements of the “Recovery” phase, however, we observed no significant difference between the two stimulation conditions (Figure 1B, Table 2). We conclude based on these data that although pH may be a factor in mobilizing intracellular Zn^2+^ upon stimulation of dissociated hippocampal neurons, different stimulation methods induce different Zn^2+^ responses through an alternative mechanism.

**Figure 4.**
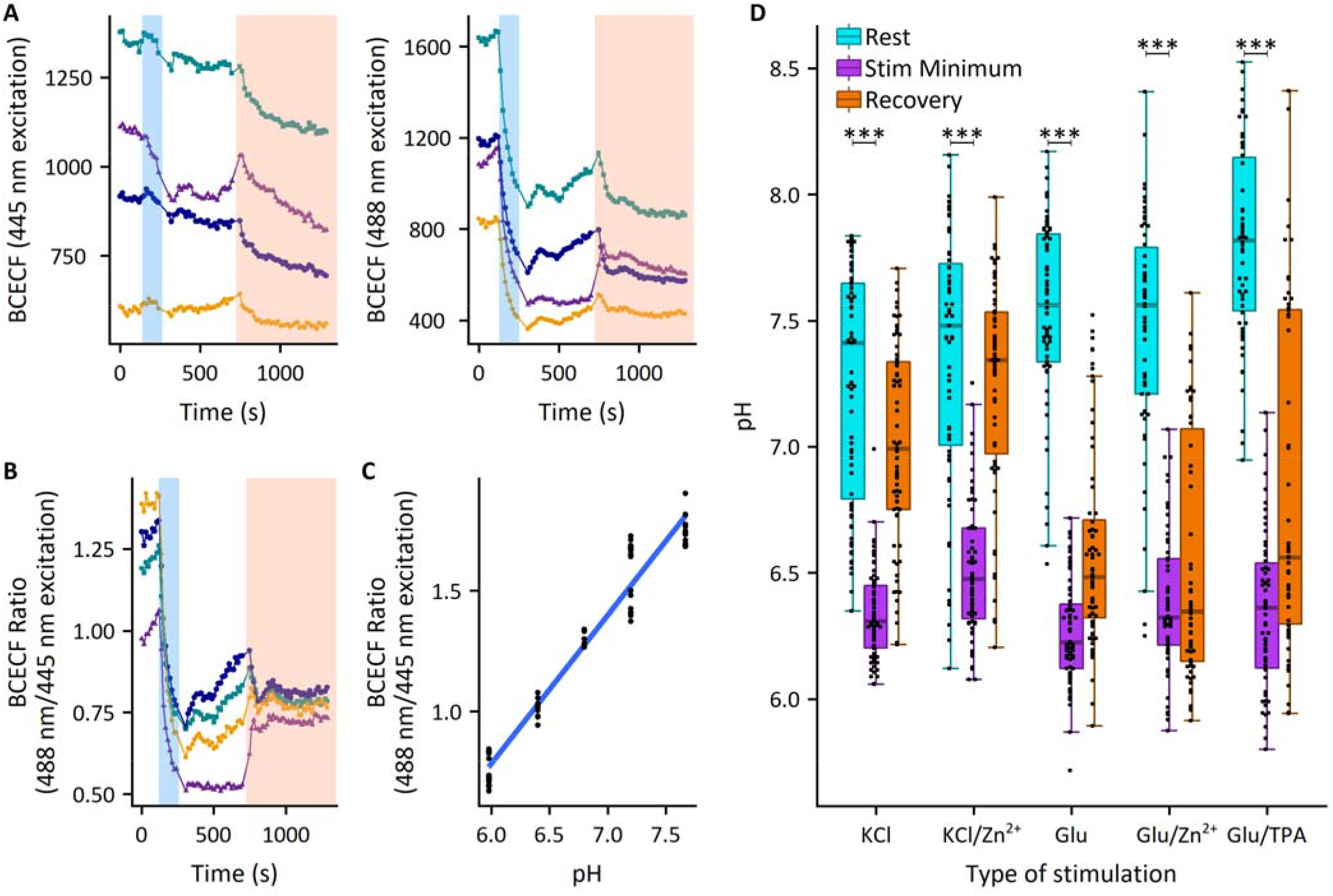
Measurement of stimulation-dependent intracellular pH responses with BCECF AM. (A) Example traces of fluorescence detected upon excitation with a 488 nm (left) or 445 nm (right) laser. Each individual trace represents the average intensity within a different cell in a single field of view. Experiments consisted of stimulation (blue box), followed by a period of recovery before addition of buffered pH media/10 µM nigericin (orange box) for calibration. Calibration pH media of the example is at pH 6.4. (B) Ratio of signals (488/445) from the same cells as in part A. Note the ratios converge upon pH 6.4/nigericin treatment. (C) Calibration curve relating BCECF ratio to equilibrated pH, obtained on a single day of experiments. Each dot represents one cell in a field of view, and 1-2 experiments comprise each pH point. (D) Box/dotplot of pH across stimulation conditions, as calculated from each individual cell BCECF ratio via the calibration curve obtained on the day of that experiment. Each dot represents values obtained from an ROI in a single cell. Rest values represent the average pH before stimulation, stimulation minimum values represent a 3-frame average around the minimal pH obtained during or directly after stimulation, and recovery values represent a 5-frame average pH before addition of pH buffer for calibration. Sample sizes: KCl = 71 cells from 7 separate experiments; KCl/Zn^2+^ = 63 cells from 7 separate experiments; Glu = 68 cells from 8 separate experiments; Glu/Zn^2+^ = 58 cells from 6 separate experiments; Glu/TPA = 58 cells from 7 separate experiments. Significance between resting and stimulation minimum values was assessed with a two-sided Wilcox Signed Rank test for paired data with all data points in each condition, ***p < 10^−10^ (KCl: p=3.3e-13, KCl/Zn^2+^: p=1.4e-11, Glu: p=7.8e-13, Glu/Zn^2+^: p=3.8e-11, Glu/TPA: p=3.6e-11).

One possible Ca^2+^-dependent mechanism that may be differentially regulated during glutamate or KCl stimulation of neurons is the generation of reactive oxygen/nitrogen species. Glutamate stimulation is known to prompt ROS production in a Ca^2+^-dependent manner^26^, and ROS are known to mobilize Zn^2+^ from cytosolic metallothioneins^27–29^, although whether the minimum timescale of this mobilization matches our observations is unclear. Some research has shown that ROS may be involved in physiological responses^30^, so ROS-dependent Zn^2+^ mobilization may not necessarily be indicative of oxidative stress. Furthermore, 60 mM KCl has been shown to not produce intracellular ROS^27^, thus potentially explaining the limited extent of the Zn^2+^ signal we observed upon KCl stimulation. Further study of ROS generation in neurons cultures upon stimulation will likely clarify the different observed Zn^2+^ dynamics.

In this study, we examined how different methods of stimulation of dissociated hippocampal neurons elicited diverse Zn^2+^ responses. We found that KCl and glutamate stimulation generated different intracellular Zn^2+^ and Ca^2+^ dynamics, and that despite literature suggesting that pH is a driving factor in glutamate-induced intracellular Zn^2+^ mobilization, the magnitudes of these differential Zn^2+^ responses failed to correlate with the extent of pH drop observed. There is thus another process necessary for further Zn^2+^ mobilization in glutamate-stimulated neurons.

## METHODS

### Neuron isolation/culture

Glass slides for imaging were coated overnight with 1□mg/mL poly-D-lysine hydrobromide in 15□mM sodium borate. Slides were washed thoroughly and coated in 50□µM iMatrix-511 (Clontech) until neuron plating.

E18 mouse hippocampi were ordered from BrainBits, LLC. Pooled hippocampi were washed with digestion medium (1X HBSS, 10□mM HEPES, 5□µg/mL gentamicin, pH 7.2) and digested 30□min in digestion medium containing 20□U/mL papain (Worthington). Samples were then washed with plating medium (MEM, 5% FBS, 0.6% wt/vol glucose) and dissociated by passing 5–10 times through full-diameter, then half-diameter flame-polished Pasteur pipets. Cells were plated on treated slides at a density of 20,000 cells/cm^2^ and were fed 3–4□hours after plating with glial-conditioned neuron culture medium (Neurobasal Medium, 2% B27 supplement, 0.3x GlutaMAX, all obtained from Thermo Fisher), and ½ media was replaced on day in vitro (DIV) 3, DIV 6, and DIV 13. Cultures were treated with 4□µM cytosine arabinoside from DIV 3 to DIV 6 to restrict mitotic cell proliferation. Cultures were grown in a cell culture incubator at 37□°C and 5% CO_2_.

Glial cells were isolated from neonatal mouse cortical tissue obtained from Charles Hoeffer’s lab at CU Boulder. Cells were dissociated from tissue as with hippocampal samples, then plated on standard cell culture dishes. Cells were fed every 3–4 days with glial medium (DMEM, 5% FBS, 0.5% pen/strep) until confluent. To generate glial-conditioned medium, neuron culture medium was added to confluent glial cultures for one day, then filtered through a 0.20□µm filter prior to addition to neurons.

### Materials

The following fluorescent small molecule dyes were obtained from Thermo Fisher: FluoZin-3 AM (#F24195), 2′,7′-bis-(2-carboxyethyl)-5(6)-carboxyfluorescein (BCECF) AM (#B1150), and Fluo-4 AM (#F14201). Stock solutions of all three dyes were prepared at 1 mM in DMSO.

Zn^2+^-specific chelator tris(2-pyridylmethyl)amine (TPA) (#723134), ZnCl_2_ (#211273), CaCl_2_ (#383147), L-glutamic acid (#G8415), Chelex (#C7901), ionophore 2-mercaptopyridine N-oxide (pyrithione) (#188549), and protonophore nigericin (#N7143) were purchased from Sigma-Aldrich. Ionomycin (#407950) was purchased from Millipore Sigma. Stock solutions were prepared as follows: TPA, 20 mM in DMSO; ZnCl_2_, 1 mg/mL in chelex-treated water; CaCl_2_, 1 M in chelex-treated water; glutamate, 10 mM in chelex-treated water; pyrithione, 5 mM in DMSO; nigericin, 1 mM in ethanol; ionomycin, 10 mM in DMSO.

Resting neuron imaging media (RNIM) was formulated as follows: 145 mM NaCl, 3 mM KCl, 1.5 mM CaCl_2_, 1 mM MgCl_2_, 10 mM HEPES, 10 mM glucose, pH 7.4. High-potassium neuron imaging media (KNIM) was made as a 2X K^+^ solution (51 mM NaCl, 97 mM KCl, 1.5 mM CaCl_2_, 1 mM MgCl_2_, 10 mM HEPES, 10 mM glucose, pH 7.4), which when added 1:1 to RNIM gave concentrations of 98 mM NaCl and 50 mM KCl.

### Equipment

Samples for all imaging experiments were imaged on a Nikon Ti-E spinning disc confocal microscope equipped with Nikon Elements software, Ti-E perfect focus system, Yokogawa CSU-X1 spinning disc head, Andor 888 Ultra EMCCD camera and Oko Labs enclosed environmental chamber set at 37 °C.

### General imaging conditions

For all imaging experiments, neuron cultures (DIV 10-14) were washed and incubated at room temperature in RNIM containing 5 μM FluoZin-3 AM, 5 μM Fluo-4 AM, or 1 μM BCECF AM for 20-30 minutes. Samples were washed with RNIM. Baseline measurements were obtained for 1-3 minutes. Cells were then stimulated with one of two basic methods: 1) 2-minute treatment of high K^+^ by mixing KNIM 1:1 with the RNIM already present, 2) 2-minute treatment with 50 µM glutamate, made at 2X in RNIM and mixed 1:1 with existing RNIM on cells. In +Zn^2+^ experiments, a final concentration of 10 µM ZnCl_2_ was added during stimulation. In +TPA experiments, a final concentration of 10 µM TPA was added 1-2 minutes before stimulation and maintained through stimulation. In +ZX1 experiments, a final concentration of 20 µM ZX1 was added 1-2 minutes before and maintained through stimulation.

### Stimulation-induced Zn^2+^ measurements with FluoZin-3 AM

Measurements were taken using a GFP channel (488 nm excitation, 525/50 nm emission), acquiring images with a 40X (NA 0.95) air objective at 300 ms exposure time, EM multiplier 300, 10 MHz camera readout speed, and 15% laser power.

After stimulation, cultures were washed with RNIM and measurements taken for 6-8 minutes. Calibrations were performed by adding 10 µM TPA (final concentration) for 2 minutes, then washing out with RNIM and adding 10 µM ZnCl_2_/0.5 µM pyrithione (final concentrations) until several minutes after a maximum signal was observed.

### Stimulation-induced pH measurements with BCECF AM

Measurements were taken using a modified GFP channel (488 nm excitation, 2% laser power, 525-542 nm emission) and a CFP/YFP 445 ex channel (445 nm excitation, 4% laser power, 540/30 nm emission), acquiring images with a 40X (NA 0.95) air objective at 300 ms exposure time, EM multiplier 300, 10 MHz camera readout speed (for each channel).

After stimulation, cultures were washed with RNIM and measurements taken for 6-8 minutes. After each experiment, media was replaced by RNIM/10 µM nigericin buffered at a different pH, which equilibrated the intracellular and extracellular pH at a specific value. On a given day, cells in 1-2 different dishes were measured at each pH, then all same-day data assembled to generate a relationship between ratio and pH. Buffered pH solutions were measured precisely each day, but generally had values around pH 6.1, 6.4, 6.8, 7.3, and 7.7.

### Stimulation-induced Ca^2+^ measurements with Fluo-4 AM

Measurements were taken using a GFP channel (488 nm excitation, 525/50 nm emission), acquiring images with a 40X (NA 0.95) air objective at 300 ms exposure time, EM multiplier 300, 10 MHz camera readout speed, and 15% laser power.

After stimulation, cultures were washed with RNIM and calibrations were performed by adding 5 mM CaCl_2_/5 µM ionomycin (final concentrations). Measurements were taken until several minutes after a maximum signal was observed.

### Image analysis

All imaging experiments were analyzed with a custom MATLAB script that imports ND2 experiment files generated by Nikon Elements software, extracts metadata, registers images, allows for manual background and cell ROI selection, and generates background-subtracted average intensity measurements.

For FluoZin-3 AM quantification, calibration data were manually inspected to obtain 3-frame average intensities around minimum (Fmin) and maximum (Fmax) values of the background-subtracted fluorescence intensity during TPA and Zn^2+^/pyrithione treatments, respectively. Fractional saturation was calculated according to the formula:

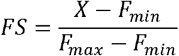

where X is the resting, peak, or recovery measurement of interest. Fractional saturation of FluoZin-3 AM was converted to an approximate intracellular Zn^2+^ concentration according to the formula:

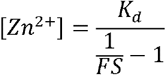

where *K*_*d*_ is the sensor dissociation constant (9.1 nM^31^) and FS is fractional saturation as defined above.

For BCECF AM quantification, BCECF ratios were calculated as (fluorescence intensity_488ex_/fluorescence intensity_445ex_). These ratios were converted to pH values according to the linear standard curve derived from pH calibration values obtained during imaging.

For Fluo-4 AM quantification, calibration data were manually inspected to obtain a 3-frame average intensity around the maximum value (F_max_) of background-subtracted fluorescence intensity during Ca^2+^/ionomycin treatment. Measurements of interest were then represented as F/F_max_. Peak excitation Ca^2+^ response was calculated in two different ways: a 3-frame average intensity around maximal peak response, and a sum of all intensities measured during the 2-minute stimulation period.

### Statistical analysis and plotting

All statistical tests were performed in R (v3.5.3) within RStudio (v1.2.1335), and are detailed in individual figure legends. For comparison of sensor responses to stimulation, two-sided Wilcox Signed Rank tests and two-sided Mann-Whitney U tests were used due to non-normality of most of the original data (as assessed by a Shapiro Wilk test).

All fits and plots were generated with R (v3.5.3) within RStudio (v1.2.1335), using the packages ggplot2 (v3.1.1), ggrepel (v0.8.0), ggpubr(v0.2), reshape2 (v1.4.3), extrafont (v0.17), dplyr(v0.8.0.1), and cowplot (v0.9.4).

## ACKNOWLEDGMENTS

We would like to acknowledge the BioFrontiers Institute Advanced Light Microscopy Core, where spinning disc confocal microscopy was performed on a Nikon Ti-E microscope supported by the BioFrontiers Institute and the Howard Hughes Medical Institute. We would also like to acknowledge the University of Colorado Biochemistry Cell Culture Core Facility for providing resources and support in culturing neurons.

## Author Contributions

L.S. and A.P. designed research and wrote the article. L.S. performed research and analyzed data.

## Funding Sources

This work was supported by an NIH Pioneer Award to A.E.P. (GM114863), a Signaling and Cell Cycle Training Grant to L.S. (T32 GM008759), and a traineeship in the IQ Biology program of the BioFrontiers Institute to L.S. (NSF IGERT 1144807).

## Notes

The authors declare no competing financial interest.

---

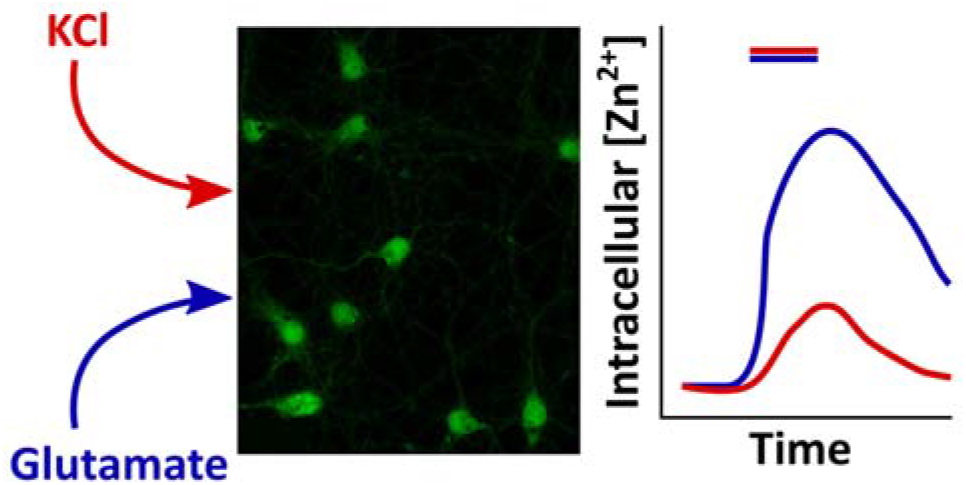

